# In depth profiling of the cancer proteome from the flowthrough of standard RNA-preparation kits for precision oncology

**DOI:** 10.1101/2023.05.12.540582

**Authors:** Filip Mundt Madsen, Annelaura Bach Nielsen, Juanjuan Wang, Josephine Kerzel Duel, Christina Westmose Yde, Martina Amnitzbøll Eriksen, Ulrik Lassen, Finn Cilius Nielsen, Kristoffer Rohrberg, Matthias Mann

**Affiliations:** Novo Nordisk Foundation Center for Protein Research; Novo Nordisk Foundation Center for Protein Research, Faculty of Health and Medical Sciences, University of Copenhagen, Copenhagen, Denmark; Center of Genomic Medicine, Copenhagen University Hospital, Copenhagen, Denmark; Department of Oncology, Copenhagen University Hospital (Rigshospitalet), Copenhagen, Denmark, and Faculty of Health and Medical Sciences, University of Copenhagen, Copenhagen, Denmark; Department of Oncology, Phase 1 Unit, Rigshospitalet, Blegdamsvej 9, Copenhagen, Denmark, and DCCC Brain Tumor Center, Department of Oncology, Copenhagen University Hospital Rigshospitalet, Copenhagen, Denmark; Department of Oncology, Phase 1 Unit, Rigshospitalet, Blegdamsvej 9, Copenhagen, Denmark; Novo Nordisk Foundation Center for Protein Research, Faculty of Health and Medical Scien1ces, University of Copenhagen, Copenhagen, Denmark, and Max Planck Institute of Biochemistry, Proteomics and Signal Transduction, Martinsried, Germany

## Abstract

Cancer is a highly heterogeneous disease, even within the same patient. Biopsies taken from different regions of a tumor may stand in stark molecular contrast to each other. Therefore, the ability to generate meaningful data from multiple platforms using the same biopsy is crucial for translating multi-omics characterizations into the clinic. However, it is generally a cumbersome and lengthy procedure to generate DNA, RNA and protein material from the same biopsy. The Qiagen AllPrep kit is an accessible, straightforward, and widely used kit in clinics worldwide to process biopsies and generate genomic and transcriptomic data from tumors. We aimed to determine if high-quality proteomics data could also be obtained from the remaining material. Here, we investigated procedures for generating deep and quantitatively accurate proteomic information in high throughput from Qiagen AllPrep flowthroughs. With a number of refinements, we obtain in excess of 10,000 quantified proteins, from 60 samples per day, achieving a substantial coverage of the total proteome. Additionally, we successfully characterize the tumors using phosphoproteomics. Combining a standard kit with in-depth proteomics will be an attractive approach for clinics seeking to implement multi-omics-based precision oncology.

## Introduction

Cancer is a devastating disease initiated by genomic damage or instability. To better understand this disease, priority has traditionally been given to the sequencing of genes and transcripts, which has increased our knowledge of the disease tremendously. Today, many leading hospitals around the world are sequencing cancer patients’ genomes and transcriptomes, aiming to match aberrations found in nucleic acid with targeted drug treatments, i.e., *precision oncology*. Regrettably, these endeavors have not yielded the anticipated substantial improvements in patient survival, particularly in the context of metastatic disease [1, 2]. To enhance precision oncology, a logical step is to characterize the proteomes of these patients, in concert with genomics and transcriptomic analyses. Proteins are the workhorses of all biological systems, healthy as well as diseased [3], and the inclusion of the proteome should add a functional layer to better characterize cellular state and signaling [4–6]. The field of mass spectrometry (MS)-based proteomics has now matured to a state that makes it attractive in clinical, translational research as well as in patient care [7–11]. Most of the material used for proteomics in the clinic is extracted from formalin-fixed paraffin-embedded (FFPE) sections, which can now be analyzed robustly and sensitively by proteomics [12]. Formalin-fixed paraffin-embedded material is routinely generated from tumor biopsies in the standard histopathological workup and used for diagnostics and patient stratification. However, the biopsy for proteomic analysis is generally not the same as the tumor material resected for genomic and transcriptomic analyses. This presents an issue for a disease such as cancer, where the different physical parts of the tumor might harbor very different genomic alterations and intracellular signaling transduction states [13, 14]. It can be a challenge to compare the various omics datasets, especially, when the goal is to understand if various genomic aberrations or transcriptomic splice forms are translated to protein-level [4].

At the Phase I Unit at Rigshospitalet, Denmark’s largest hospital, patients with metastatic cancers and exhausted treatment options are offered enrollment into a precision oncology trial termed CoPPO [2]. After consent, they have their metastatic lesions biopsied for genomic and transcriptomic sequencing at a scale of several hundreds of patients per year. The sequencing data is included in an integrated clinical report together with the histopathological evaluation (based on an adjacent FFPE biopsy) and supporting clinical decision making at the National Molecular Tumor Boards, with the aim to guide further patient-specific care. The Qiagen AllPrep kit, widely used for extracting RNA and DNA, leaves behind a protein-rich flowthrough. This flowthrough represents an attractive resource for proteomics analysis. The Phase 1 Unit has prospectively collected thousands of these flowthroughs from sequenced patients, stored at-80°C.

Here, we report on our efforts to analyze the saved flowthrough fractions, referred to herein as the *protein-fractions*, using MS-based proteomics. We aim to explore the feasibility of generating high-quality proteomics data from these protein-fractions, the potential to extract meaningful biological and clinical information, and scale up the sample throughput. Additionally, we focus on the enrichment of phosphopeptides. We believe that utilizing these protein-fractions will bring us one step closer to the inclusion of proteogenomic-based precision oncology in Molecular Tumor Boards.

## Results

### Obtaining meaningful proteomics data from the protein-fractions

For a more detailed description of our work, please see the *Materials and Methods* section of the paper. When using the Qiagen AllPrep kit, biopsies are denatured using guanidine-HCl, which is used in many proteomics workflows already. The ensuing denaturing milieu will inhibit enzymatic activity and should keep the proteome intact during and after DNA and RNA extraction. The flowthroughs that are left after DNA and RNA extraction, contain proteins that can be precipitated at room temperature using the provided Advanced Protein Purification buffer (APP; Qiagen). These protein-fractions (flowthroughs) then also contain guanidine-hydrochloride, dithiothreitol (DTT) and a varying amount of ethanol. Unfortunately, the final concentrations and compositions are hard to determine. The protein pellet obtained after the addition of APP can be analyzed using techniques such as sodium dodecyl sulfate (SDS)-page, Western-blotting and, allegedly, mass spectrometry. We do observe protein bands when the pellet is re-suspended in SDS, or 2-amino-2-(hydroxymethyl)propane-1,3-diol;hydrochloride (tris-HCl; pH 8.0) buffers, sonicated and loaded on a gel for Coomassie Brilliant Blue staining. Several ways to re-suspend and clean up the proteins, and de-salt the resulting peptides, after enzymatic digestion and before mass spectrometry analyses are depicted in **Figure 1a**. Initially, we attempted to re-suspend the proteins by simply adding the provided APP buffer, which resulted in a large insoluble pellet. We then dried the pellet and re-suspended it in either sodium deoxycholate (SDC; PreOmics) or Tris-HCl buffers with the aid of sonication (Bioruptor® Plus) and vigorous pipetting. The resulting solution, from either buffer, was digested using Lys-C and trypsin, and after acidification the peptides were de-salted on styrene-divinylbenzene reverse-phase resin (SDB-RP) StageTips [15] or alternatively solid-phase extraction (Sep-Pak®) C_18_ columns. Nano-Drop concentration determinations showed well-defined protein curves at the 280 nm, and concentrations ranging from 0.1 µg/µl to 3.0 µg/µl. However, despite our best efforts, the peptides obtained from the APP precipitation and re-suspension in any of the above buffers did not yield satisfactory results. The spectra obtained with an Easy-nano-liquid chromatographer (nLC; Agilent) coupled to an Orbitrap mass spectrometer (Exploris 40™; Thermo Fisher Scientific) setup were of poor quality compared to those obtained from quality control HeLa injections (**Figure 1b**, top and middle panels, respectively) and resulted in the quantification of only 500 proteins. Next, we digested and de-salted the samples using the Protifi S-Trap™ kit (Protifi) which resulted in better spectra and doubled protein identification to roughly a thousand quantified proteins. Note that the guanidine-HCl buffer is not compatible with the initial SDS-containing Protifi buffer (it turns into a gel-like substance). As a result, the first step of the Protifi protocol involving the use of the SDS-containing buffer was omitted. Finally, we precipitated the proteins using ice cold acetone overnight. The resulting protein pellet was resuspended in SDC, tris-HCl (pH 8.0), SDS 0.1 to 4 %, or 8M urea buffer together with sonication to better solubilize the pellet. The proteins were then reduced and alkylated using DTT and iodoacetamide and digested using LysC for 2 hours and then with trypsin overnight. The resulting digest was cleaned on either SDB-RP StageTips or SepPak columns of various sizes. We obtained the best yields with the SepPak columns. The resulting desalted peptides generated much richer spectra (**Figure 1b**, bottom panel) enabling quantification of more than 3,000 protein groups from the first test samples.

**Figure 1.**
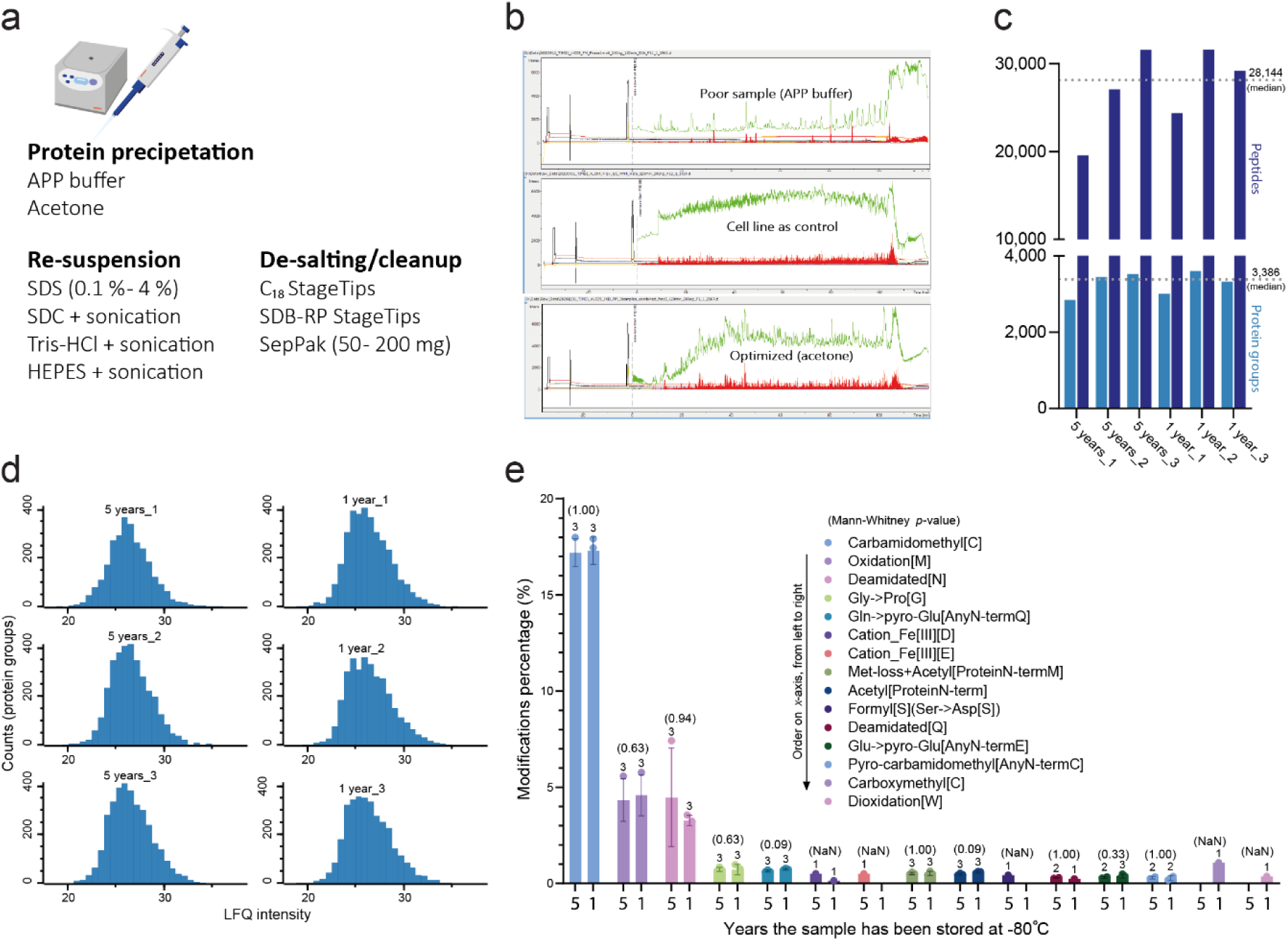
Clean-up and mass spectrometry acquisition of full proteomes from the flow through-fractions. **a)** Several alternative processing workflows to obtain clean peptides for MS analysis. **b)** Chromatograms that resulted from peptides resuspended after APP precipitation, (top) an HeLa quality sample (middle), or peptides processed after acetone precipitation (bottom). **c)** Number of peptides and protein groups quantified after acetone precipitation and digestion of six proteins fractions after various storage times. **d)** The distribution of the proteins in panel “c” for all six samples, after log_2_ transformation. **e)** An open search (pFind) identified several post-translational modifications on the peptides in these six samples. None of the modifications were significantly different in samples stored for one year, versus 5 years at-80° C (Mann-Whitney U test *p*-value). The amino acids that were modified are denoted with their single letter or three letter symbols (e.g., “[S]”, and “Ser” both denote serine).

After sequential DNA and RNA extraction, the Qiagen AllPrep kit yielded a flowthrough of approximately 800 µL volume that contained the associated proteins. From this we processed 250 µL for acetone precipitation. Given the difficulty of estimating the final composition of the lysis buffer that we received our protein-fractions in, we decided against upfront protein concentration determinations. The presence of DTT invalidates the use of a standard Pierce™ BCA Protein Assay Kit (Thermo Fisher Scientific) and the Nano-Drop absorption curves at 280 nm indicate interfering contamination. Instead, we back-calculated the theoretical initial protein amount assuming a 50 % material loss from the collection of proteins to after de-salting of the peptides. This results in about 0.2 µg/µl concentrations and about 50 µg of total protein for the entire 250 µL protein-fraction. However, these concentrations varied substantially from sample to sample and are likely tissue specific. In any case, when desalting only 10% of the total peptides after protein digestion, we obtained sufficient amounts of clean peptides to quantify the entire proteome multiple times. This indicated that protein amount is not a limiting factor in these samples.

After determining the proteome preparation procedure, we proceeded to analyze six patients with different metastatic lesions from various cancer types. These were selected to represent different historical storage times: three samples had been stored for 5 years and three for one year. After acetone precipitation and SepPak clean-up, we quantified a median of around 28,000 precursor peptides, mapping about 3,300 protein groups using a single-run data-dependent acquisition (DDA) approach on an Easy-nLC Exploris setup, with 145-minute gradients and searched using MaxQuant [16] (**Figure 1c**). These samples show expected and comparable distributions of their proteins (**Figure 1d**). Importantly, when performing an open database search using the pFind software [17] (v. 3.0), to broadly assess various post-translational modifications, we did not see any difference in their relative abundance between storage for 5 or one years at-80° C. The most common post-translational modification was carbamidomethylation of cysteine (17.2 % if stored for 5 years and 17.4 % if stored for one year), which is a modification introduced on purpose during reduction and alkylation. The second most common modification was oxidation of methionine (4.3 vs. 4.6%), followed by deamidation of the N-terminus (4.5 vs 3.3%). The full list of detected post-translational modifications can be seen in **Figure 1e**. Our results indicate that the integrity of these cancer proteomes is maintained at least up to five years of storage, opening the possibility for retrospective analyses of these cohorts.

### Obtaining meaningful biological and clinical data from the protein-fractions

The next step was to evaluate if we could extract any biological signal from our proteome measurements. The initial six samples that we had analyzed for their post-translational modifications were collected from various primary cancers such as small intestine adenocarcinoma, tonsillar squamous cell carcinoma and thymoma, and from different metastatic sites, including liver, lymph node, and kidney tissue. For three of these tested samples, we had information on both the cancer of origin, and which tissue the metastatic biopsies were resected from. For the three samples with known primary and metastatic sites, we performed a single-sample gene-set enrichment analysis (ssGSEA) using cell type signature gene sets from the MSigDB (v. 7.4) C8 database (http://software.broadinstitute.org/gsea/msigdb/). This database has “gene sets that contain curated cluster markers for cell types identified in single-cell sequencing studies of human tissue”, which allowed us to identify significantly enriched cell type signatures. In this analysis we matched the tissue from the cancer of origin in one sample with the tissue of the metastatic sites in the remaining two (**Figure 2a**). Note that for the two samples where the metastatic site was identified, the cell types corresponding to the cancer of origin were not present in the MSigDB C8 database (specifically for the tonsillar squamous cell carcinoma and thymoma type B2), hindering this analysis. Still, the results indicate that the proteomes of these samples retain quantifiable cell type specific information, even when stored for up to five years. This paves the way for retrospective identification of primary and metastatic sites in these samples. Furthermore, our proteomic findings quantified numerous well-established oncogenes and “cancer markers,” such as EGFR, IDH1, IDH2, AKT1, AKT2, MAPK1, and MAPK3, carrying potentially valuable clinical insights (**Figure 2b**).

**Figure 2.**
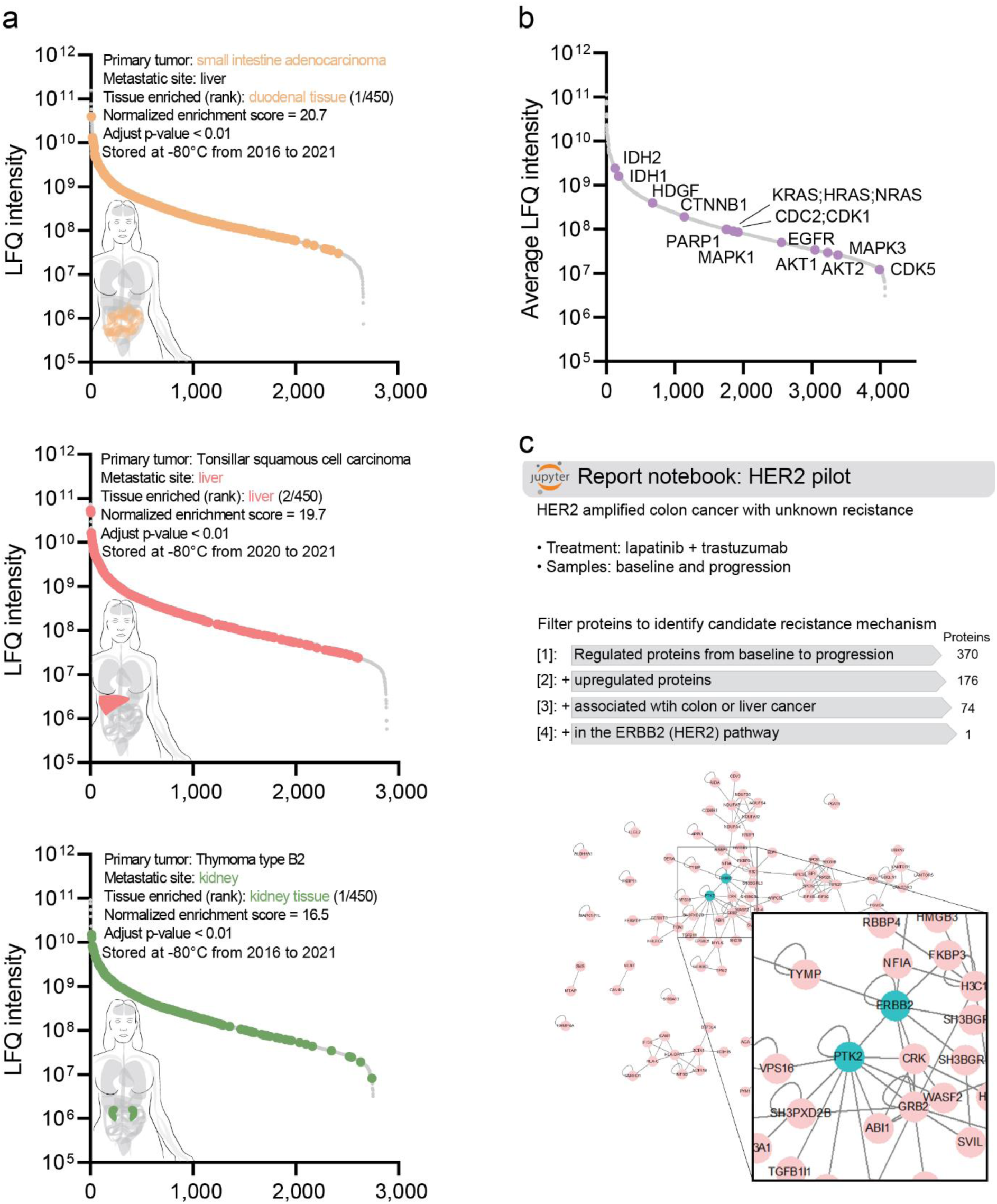
Quantification of tissue specific markers and molecular correlates of a treatment resistance. **a)** Using the MSigDB C8 cell type signature gene sets, a gene-set enrichment on a single sample basis correctly identified either the origin of cancer or the site of the metastatic lesion for all three analyzed patients (left side). LFQ = label free quantification. Colored dots are genes that belong to the noted gene set, and in the end results in the normalized enrichment score for each tissue. **b)** Several important cancer markers were quantified from these protein-fractions (purple dots). The LFQ intensities are averaged from all proteins quantified in at least one of the six samples illustrated in Figure 1, panel “d”. **c)** In the case of a patient with HER2 amplified colon cancer and unknown resistance, our data identify a possible culprit: PTK2/ FAK1. These findings are based on the Clinical Knowledge Graph and are displayed as a Jupyter notebook excerpt. The network shows all proteins remaining after the second to last filter (associated with colon or liver cancer; n = 74) while highlighting PTK2 and ERBB2 (Her-2-neu) in turquoise.

To further evaluate the clinical utility of analyzing the proteomes of these protein-fractions, we processed two metastatic lesions from a patient with an ERBB2 (Her2-Neu) amplified colon cancer that had metastasized to the liver. This patient had failed anti-Her2-Neu treatment with trastuzumab and anti-EGFR treatment with lapatinib. However, upon recurrence the patient had not developed any new genomic alterations that could explain the resistance. When analyzing the protein-fractions from both baseline and progression metastatic biopsies, we quantified more than 3,700 proteins in single-run DDA mode. We focused on the top 10 % of proteins that changed the most in their relative abundance from baseline to progression, and only those that were upregulated because they are potential therapeutic targets. Using the Clinical Knowledge Graph, a bioinformatic framework developed in our laboratory [18], we filtered for proteins associated with colon or liver cancer, AND with trastuzumab and lapatinib AND with Her2-Neu signaling. This highlighted one protein as a possible reason for resistance to treatment, namely focal adhesion protein kinase 1 (PTK2/FAK1; **Figure 2c**). This kinase was here identified solely in our proteomics data and has been described in the literature to be associated with trastuzumab resistance in ERBB2 amplified breast cancers [19]. This is clearly an interesting finding in this context. It also underscores the ability to obtain additional clinical information from these flow through protein-fractions and strengthens the rationality of our approach.

### Increasing proteomic depth

Single-run analysis in a clinical setting reaching a depth of below 4,000 proteins is already quite useful but is far from the complete proteome. To increase proteomic depth, we decided to analyze a set of 27 flow-through protein-fractions from BRAF-mutated (V600E) solid tumors using a different strategy. These samples are part of a larger cohort (n = 46) that we analyzed in one batch and deposited together online (see section *Data availability*). This collection represents a clinically relevant cohort with different primary tumors and metastatic sites (**Figure 3a**), as well as several types of biopsies, sometimes serial biopsies from the same patient, at “baseline” (n = 16), “new baseline” (n = 2), “on treatment” (n = 7), “progression” (n = 1) and “new genomic biopsy” (n = 1). To increase proteomics depth, we decided to analyze these protein-fractions in data-independent acquisition (DIA or dia) mode, instead of data-dependent mode (DDA), which results in higher data completeness [20, 21]. Both strategies were tested on an Easy-nLC coupled to a timsTOF Pro (Bruker) with Parallel Accumulation Serial Fragmentation (PASEF; ddaPASEF and diaPASEF, respectively) [22–24]. With the advances in DIA search algorithms, we expected to increase our proteomics depth and decrease missing values. Indeed in 120-minute gradients, we quantified a median of 4,351 protein groups using ddaPASEF followed by MaxQuant database search. For diaPASEF with DIA-NN search [25] this increased to 9,173 protein groups. Thus, use of DIA instead of DDA more than doubled the number of quantified proteins for our samples. Additionally, missing values decreased from 42 % to 20 %. We therefore continued with diaPASEF and DIA-NN (**Figure 3b**).

**Figure 3.**
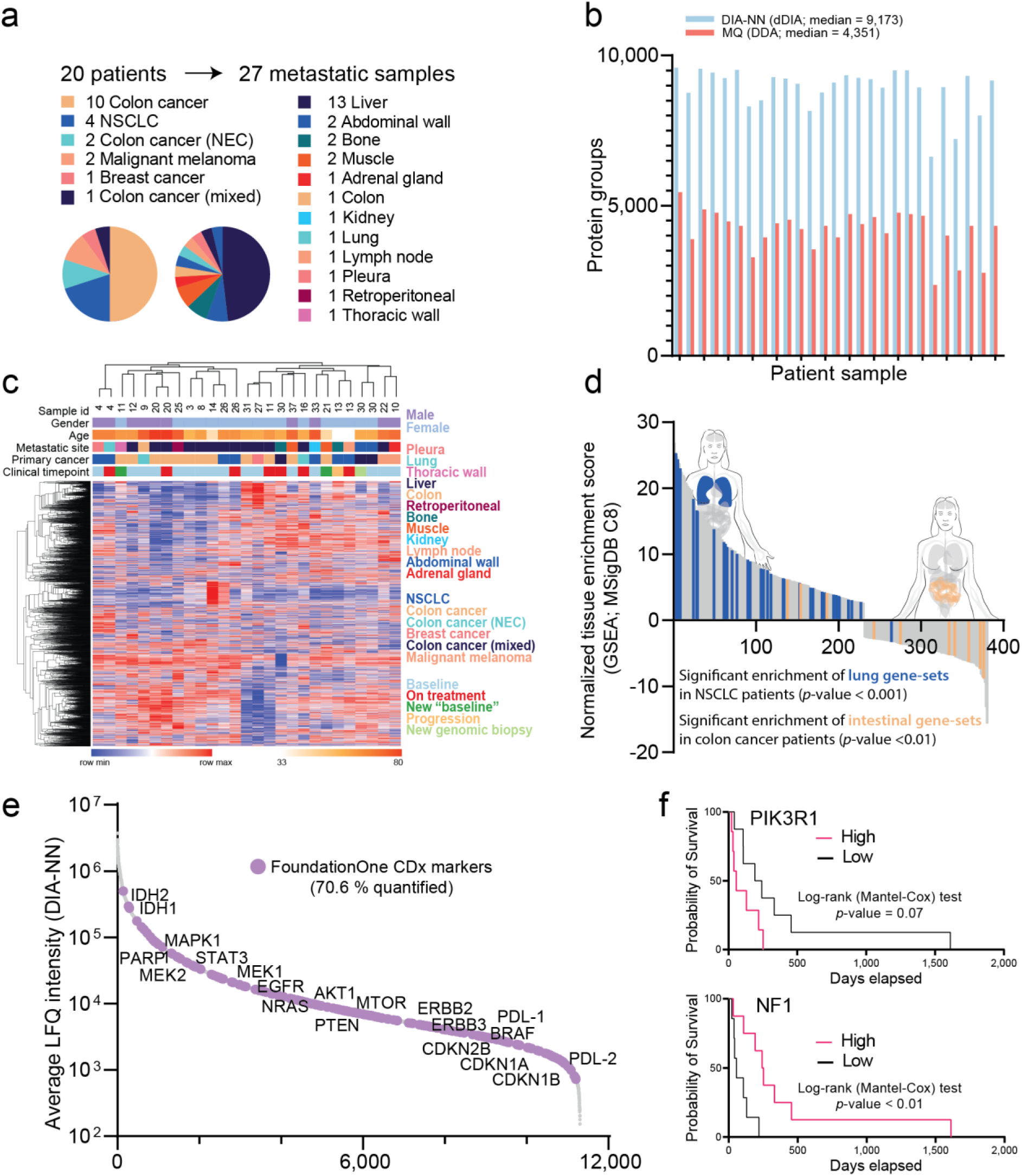
Increasing depth of the proteome reveals pathophysiology and clinical markers. **a)** Twenty-seven protein-fractions from 20 patients were acetone precipitated and prepared for proteome analysis. All biopsies were collected from a metastatic lesion in the patient. **b)** Moving from data-dependent acquisition (DDA) to data-independent acquisition led to an increase in quantified proteins. Data acquired in DDA mode was searched using MaxQuant, while DIA data was searched using DIA-NN with on-the-fly generation of a spectral library for re-annotation. **c)** Unsupervised, one minus Pearson correlation hierarchal clustering of all samples reflects underlying biology, with samples from the same patient clustering together, colon cancers and liver metastases forming distinct sub-groups. Clustering of samples in the heatmap is based on complete protein values, corresponding to 4,688 protein groups. “Sample id” are specific numbers for one patient, i.e., all with number “13” are from the same patient, but at different time points. **d)** In cell type signature gene-set enrichment, we identified significant intestinal gene-sets in the colon cancer samples (n = 10) and lung gene-sets in the NSCLC samples (n = 4). Only significantly enriched gene-sets are shown. Adult lung tissue gene-sets are shown in blue, adult intestinal tissue gene-sets are yellow, and all other tissue gene-sets are in grey. *P*-values were calculated using the chi-square tests. **e)** Overlay of the FoundationOne®CDx gene-list (Roche) onto our proteomics data. More than 70 % of the FoundationOne genes are translated to proteins and quantified in our analyses. **f)** Possible prognostic markers from analyzing the overall survival of the “baseline” samples.

The 27 BRAF-mutated samples were collected over a 5-year period and the numbers of quantified peptides and protein groups were similar across all years (**Supplementary** Figure 1a). Unsupervised clustering of the samples reveals underlying biology in the form of colon cancers and liver metastases loosely forming larger groups (**Figure 3c** and **Supplementary** Figure 1b) and paired most of the serial biopsies from the same patient, see for example “on treatment” and “progression” biopsies from patient with “sample id” 13; **Figure 3d**. When enriching the two largest groups of cancers of origin colon cancer and non-small cell lung cancer (NSCLC), the cell type signature specific gene-sets revealed a highly significant enrichment of *adult lung tissue gene-sets* in the NSCLC group (*p*-value < 0.001), and *adult intestinal tissue gene-sets* in the colon cancer group (*p*-value < 0.01; **Figure 3d**).

To evaluate the predictive nature of these data, we overlayed the FoundationOne ®CDx gene-list (Roche) onto our proteomics data. This list of genes is highly curated by experts across all branches of oncology and compiled to provide physicians with clinically actionable information, including both predictive markers and those carrying evidence of therapy resistance [26]. Overall, our proteomics data quantified the corresponding proteins of 71 % (222 out of 315) of these genes (**Figure 3e**). Proteomics covered the majority of these targetable proteins with high intensities in all “on treatment” and “progression” samples analyzed, which is information that we believe clinicians would value greatly. Furthermore, some of these markers seem to carry prognostic information as well. Analyzing the baseline samples revealed prognostic value of the NF1 protein (*p*-value < 0.01), where high levels seem protective for patients, while high levels of PIK3R1 indicated a worse prognosis (*p*-value = 0.07; **Figure 3f**).

### Increasing sample throughput

In order to scale up and increase our sample throughput, we evaluated the trade-off between throughput and proteomics depth when decreasing the gradient time from 120 minutes to 44 minutes using an Evosep liquid chromatography system, which is designed for robust clinical applications [27]. This approach should make our method more attractive in a clinical setting, where large numbers of samples need to be analyzed. This reduction in gradient time corresponds to a throughput of 30 samples per day (SPD). To check if this 2.5-fold increase in throughput would retain accurate quantitation of the proteome, and more importantly, retain biological and clinical information, we re-analyzed the 27 BRAF-mutated protein fractions using the remaining clean peptides. We acquire all proteomes within 24 hours, in diaPASEF acquisition mode, using a timsTOF SCP mass spectrometer and analyzed as before. With this setup, we reached a median quantified proteome depth of 7,607 protein groups (**Figure 4a**), which corresponds to 82 % of the previous 120-minute gradient acquisition. There was a very high correlation of protein groups quantified between the 120-minute and 44-minute gradients (Pearson *r* sq. = 0.97; **Figure 4b**), as well as a high average correlation of protein group intensities within samples (average intra Pearson correlation 0.95 with a standard deviation of 0.01). Consequently, an unsupervised clustering of the samples indicated that biological, and by extension, clinical information was retained with a remarkably similar paring of samples (**Figure 4c**).

**Figure 4.**
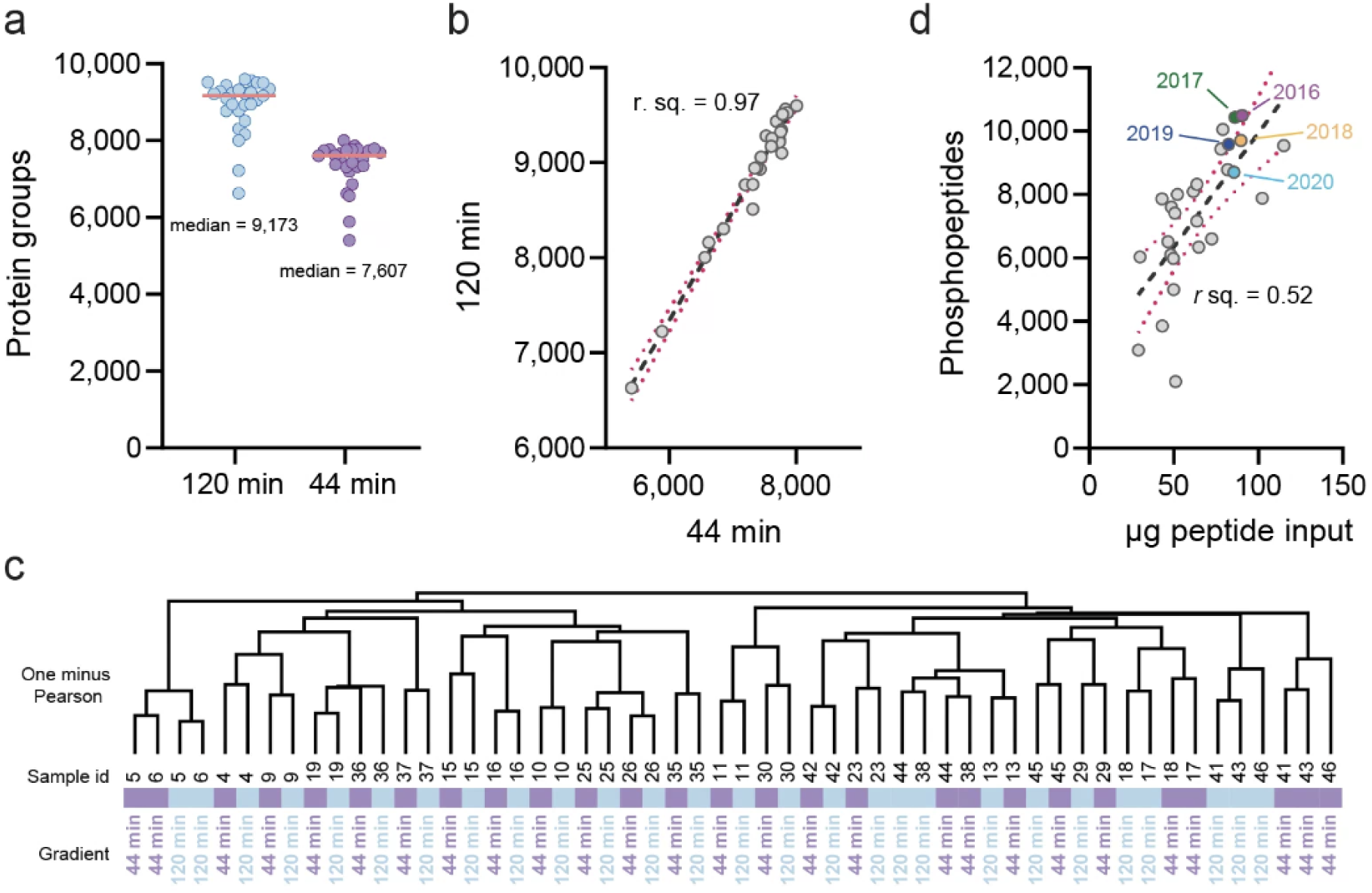
Increased throughput with retained proteomic depth and biological signatures and phosphoproteome analysis. **a)** Reducing the chromatographic gradient from 120-minute to 44-minute only reduced quantified protein groups slightly. This more economical setup with increased throughput from less than 12 samples per day (SPD) to more than 30 SPD. **b)** Samples with many protein groups in the 120-minute gradient also yielded the most proteins groups with the 44-minute gradient (30 SPD) Evosep gradient (Pearson *r* sq. = 0.97). **c)** Any given sample showed high correlation between gradient lengths. The same samples cluster together in almost all cases, when analyzed with an unsupervised, one minus Pearson hierarchical clustering. In the cases where they do not, the same cluster of samples line up perfectly. **d)** Phospho-site quantification scale with input (µg on the x-axis, ranging from 29 to 115 µg input) peptide material for the immobilized metal affinity chromatography enrichment. This levels off at around 75 to 100 µg, with a Pearson correlation of 0.52. Years of sample storage does not seem to influence the number of phosphopeptides enriched. One sample from each storage year is color coded, matched with similar input material used for enrichment. The dashed black lines and the red dotted lines in panels **b)** and **d)**, denote the means of the linear regression and the errors (95 % confidence bands), respectively.

### Measuring the phosphoproteome in the protein-fractions

The phosphorylation status of proteins is a key aspect of cancer-related pathways and can provide valuable insights into the progression of the disease [3]. As a final step, we therefore aimed to enhance our understanding of intracellular cancer signaling by enriching and quantifying phosphopeptides. To this end, we used the left-over material of peptides from the BRAF-mutated samples for phosphopeptide-enrichment. Material input ranged from 29 µg to 115 µg, and enrichment was done on an AssayMAP robot using immobilized metal affinity chromatography (IMAC) high-capacity Agilent AssayMAP Fe(III)-nitrilotriaceticacid (NTA) cartridges. Spectra for the phosphoproteomes were acquired on an Orbitrap mass spectrometer in DIA mode on 70-minute gradients. Searching the results using directDIA in Spectronaut (v. 14, Biognosys) quantified between 2,101, and 10,492 phosphopeptides, which correlated moderately to the peptide input for the enrichment (Pearson *r* sq. = 0.52; **Figure 4d**). As in the proteome measurements, we did not observe any obvious bias in the number of enriched phosphopeptides across storage times from 5 to one year; **Figure 4d**). The phosphoproteomics data resulted in similar clusters as for the proteomics data (**Supplementary** Figure 2), reflecting similar biology or the fact that phosphopeptide abundances to some extents are proxies to their corresponding protein abundances [7]. The total of about 20,000 phosphopeptide sites map to 4,839 genes that in turn cover 130 (41 %) of the 315 FoundationOne medicine markers discussed in relation to **Figure 3e**. This is a lower coverage than on protein level, however, with an additional dimension of information. For example, we quantified the AKT1 serine 124 phospho-*site*, which has been implicated in AKT-activation [28] and associated with pancreatic ductal adenocarcinoma [29]. We quantify multiple phosphorylation sites on APC, including a serine at residue 2449, whose phosphorylation has been associated with multiple cancers including breast, lung and gastric cancer [4, 30]. We also quantified the serine 151 residue on BRAF. This site is well-studied and has been linked to both NRAS and ERK regulation, with an effect on NRAS [31] and RAF1 signaling and altered cell growth [32], as well as the equally well-studied serine 365 site on BRAF that is regulated by AKT signaling and leads to inhibition of BRAF and altered cell growth [33, 34]. Another phospho-site that we quantified on BRAF was serine 729 that leads to BRAF activation when phosphorylated [35]. On EGFR we recorded several phosphorylation sites, including threonine 693 that is involved in ERK1/2 signaling [36], cell cycle regulation and EGFR activity [37, 38] and serine 1166 that has been associated with multiple cancers [39–41]. These findings are a small selection of our total data that suggest the increased level of detail in clinical information that can be obtained from quantifying phosphorylation sites. We conclude that one can readily enrich and quantify phosphopeptides from these protein-fractions, which cluster samples similarly to the proteome, indicating high quality, while adding another dimension of clinical information.

### Increasing proteomic coverage with the Orbitrap Astral instrument

In our pursuit of enhancing proteomic coverage, we sought to assess the potential of the Orbitrap Astral mass spectrometer from Thermo Fisher Scientific [42, 43] within our experimental framework. To achieve this, we processed a cohort of 96 samples derived from metastatic lesions using the Qiagen AllPrep kit. The samples underwent upfront transcriptomic and genomic profiling, followed by processing with the KingFisherTM Flex System (Thermo ScientificTM)30 utilizing MagReSyn® Hydroxyl beads (ReSyn Biosciences) to enable efficient batch processing on 96-well plates. Subsequently, peptides were analyzed using the Evosep LC system with a 21-minute gradient, effectively doubling our daily sample throughput from 30 to 60 (i.e., 60 SPD). This approach yielded a substantial increase in proteome depth, identifying 10,197 protein groups, representing a significant improvement over our previous endeavors – a feat made more remarkable considering the reduced gradient length (**Figure 5a**). When compared to the 30 SPD results, our throughput increased from 173 to an impressive 485 protein groups per minute. Furthermore, the median peptide counts per sample reached 125,348 (**Figure 5b**), with an average protein coverage of 12.3 peptides per protein. Moreover, this workflow also leads the quantification with a low reproducible coefficient of variation (CV) of ∼4.6%.

**Figure 5.**
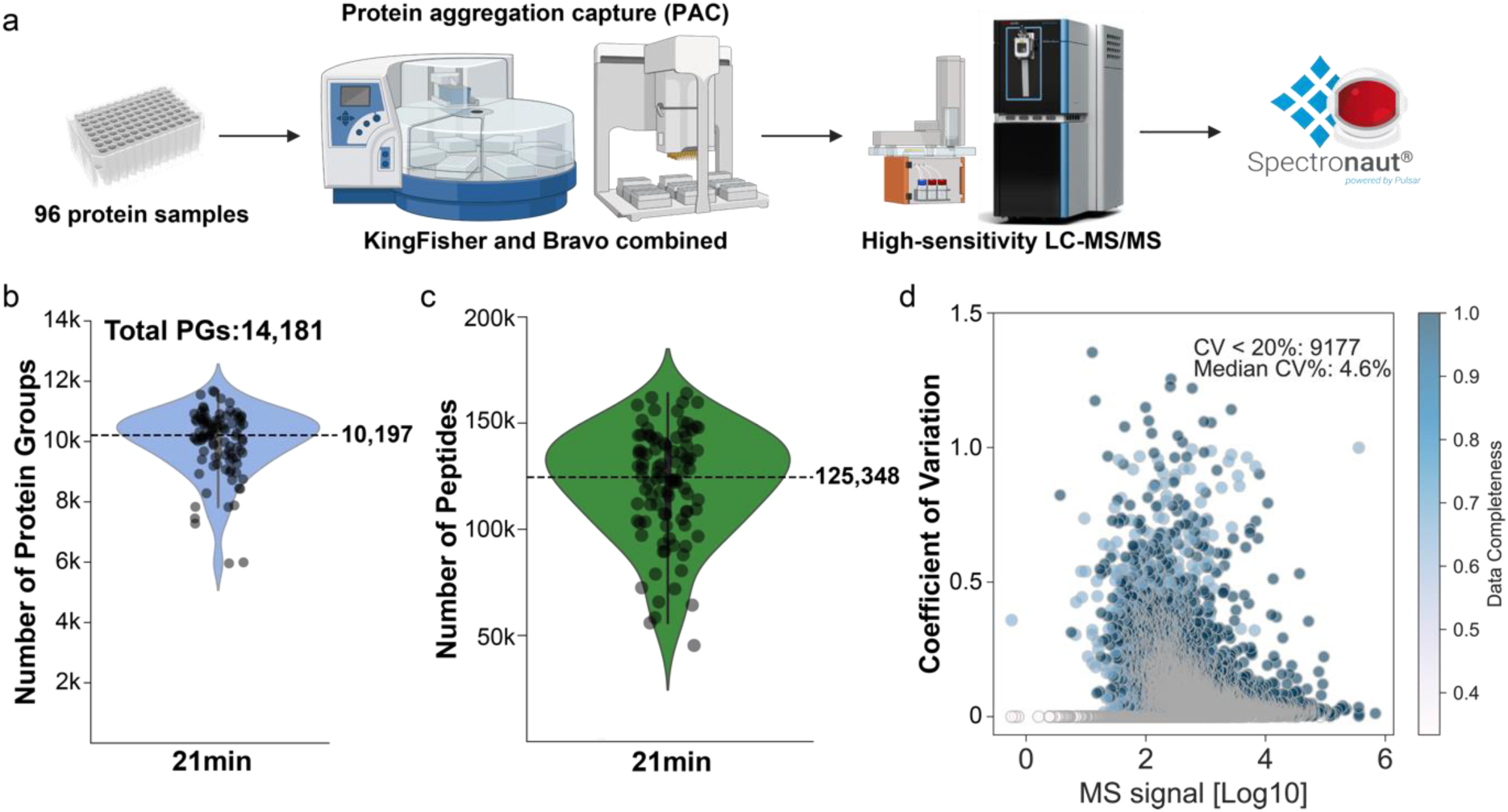
Deep proteome coverage with short gradients and Orbitrap TOF. **a)** Workflow of automated high-throughput with high-sensitive LC-MS/MS-based proteomics. Orbitrap Astral mass spectrometer coupled with Evosep with the chromatographic gradient of 21 minutes (60SPD) led to a very deep quantified proteome. **b)** and peptides numbers **c)**. Dash lines indicate the median count of protein groups and peptides**. d)** Coefficient of Variation (CV) for the 96 pooled QC samples was calculated on the raw intensity.

## Discussion

The ability to measure multiple omic-types from a single tissue biopsy is important when striving to acquire an accurate systemic view of a physiological or pathophysiological state. This is especially true for a heterogeneous disease such as cancer, where the molecular profile from a neighboring tissue area often differs substantially from the next. Correlating genomic aberrations with transcript expressions and/or protein levels becomes increasingly difficult and misleading when the distance between biopsies increases. Several studies measure DNA and mRNA from a single biopsy sample, however, few have obtained protein-level information from the exact same cancer biopsy. There are efforts to shave off FFPE sections from a single block and analyze them in a sequential manner, reaching near-same tissue location [44]. Those efforts must be performed after the biopsy has been FFPE treated and not on fresh frozen material, they are also not strictly in the same location, and laborious, requiring specialized expertise and equipment. So far, the closest to a unified analysis of all omic-types from the same biopsy has been to cryo-fracture the tissue and then distribute aliquots to the various platforms. This ensures that all technology platforms have an average view of the same piece of tissue [4, 7]. However, this, again, requires specialized machinery and can be rather laborious, especially for a large sample cohort (> 50 samples). On the other hand, it starts from fresh frozen tissue, which might be advantageous – especially if one wants to enrich phosphorylated peptides.

Conceptually, the analyses of the flowthrough from the Qiagen AllPrep kit lead to the parallel analyses of DNA, mRNA, and proteins from the exact same biopsy. The kit itself is readily available and easily used by laboratory personnel, without the need for any specialized equipment. The protein-fraction – that is the resulting flowthrough after collection of the nucleic acid fractions – is basically a free left-over, and as we show here, it is just acetone precipitation away from being mass spectrometry-ready. A note of caution is that the supplied APP buffer does not work for protein purification for MS-based proteomics and is only suitable for SDS-page electrophoresis and Western blotting.

Given the results shown here, it would be attractive to create ubiquitous repositories and biobanks with this kind of material. While FFPE blocks are available in the millions, and in most hospitals, the AllPrep kit from Qiagen is mostly used in sequencing centers, and not always with the foresight of keeping the remaining protein-fraction for future analyses. Nevertheless, when available, these kits contribute easily accessible material for multi-omic analyses, including deep proteomes, from one biopsy. Previous studies have used the Qiagen AllPrep kit for proteomics with acetone precipitation [45–49]. Herin, we report on single-shot deep proteomes from the protein-fractions remaining after using the Qiagen AllPrep kit from a tumor biopsy, and importantly we carefully characterize them for stability after years of storage time [50–52]. In this study, we open for the quantification of DNA, RNA, proteome, and phosphoproteome from the exact same biopsy in a clinical Phase 1 setting. Note that there is no principal reason to stop there. Additional *omes*, such as the lipidome, or the metabolome, or other post-translational modifications, could be acquired as well, and we encourage our colleagues to explore this. Higher-dimensional molecular characterization should lead to increased resolution of any physiological, or pathophysiological, state at hand – increasing the chances to translate such information into better patient care in this budding age of precision medicine.

We process the samples on an Agilent Bravo liquid handling platform, utilizing an LC gradient length of only 21-minutes – corresponding to 60 cancer proteomes per day, and quantifying samples on an Orbitrap Astral mass spectrometer leads to in-depth quantitative coverage. In turn this opens for ambitious proteomic projects with cohorts in their thousands, while achieving deep proteomic coverage of tumor tissues. With an increased proteomics depth comes an increased likelihood of identifying resistance mechanisms and druggable proteins. As the proteomics community steadily progresses towards such future endeavors [53], we aspire to actively contribute to the expansion of both depth and breadth in the near term.

In summary, we here produced several hundred micrograms of material from one protein-fraction and obtained high-quality deep proteomes with more than 10,000 protein groups from samples stored up to five years. Additionally, we increased the sample throughput to 60 samples per day while maintaining a high depth of their proteomes. We also demonstrated the possibility of enriching phosphoproteomes from these protein fractions. All of this can be achieved using the same biopsy that has already been used for genomic and transcriptomic analyses. The ability to obtain deep proteomes and phosphoproteomes with high throughput from the same biopsy used for genomics and transcriptomics enables a unique and ideal environment for proteogenomic analyses in the clinical setting. We hope that this will increase the prospects of successful precision oncology using more comprehensive molecular data, and the inclusion of proteomics, phosphoproteomics, and proteogenomics information in Molecular Tumor Boards.

## Acknowledgments

Our gratitude goes out to the patients who enroll in clinical trials and research studies, who enable this research, benefitting all cancer victims. We also acknowledge the staff at Genomic Medicine (Kennedy center, Glostrup, Denmark) who performed the extraction of DNA and RNA, leaving us with the left-over protein-fractions. Additionally, we thank our proteomics colleagues at the Novo Nordisk Foundation Center for Protein Research and Max Planck for competent help and feedback along the way.

## Funds

This work was supported by grants from Novo Nordisk Foundation (Grant agreement NNF14CC0001) and Danish Cancer Society (grant no. R316-A19079).

## Materials and Methods

### Patient selection and biopsy

All available samples collected in the CoPPO trial [2] were enrolled. The CoPPO trial comprises patients with advanced solid malignancies who were referred to the phase I unit at Copenhagen University Hospital, Rigshospitalet (Copenhagen, Denmark).

A fresh tumor sample primarily from metastatic sites was collected using either core-needle biopsies (18-gauge needle) or surgical resection samples under local anesthesia. Three samples were obtained from the same lesion, with two stored in RNAlater (Life Technologies) for RNA expression analyses and DNA gene mutation analyses and one sample formalin-fixed, paraffin-embedded (FFPE) for histopathologic analyses to confirm the suitability and representativeness of the material.

The study was conducted in accordance with the Declaration of Helsinki and approved by an institutional review board and the Regional Ethics Committee (Danish Ethical Committee, file number: 1300530). All patients provided signed informed consent.

### Sample preparation of the protein-fractions

The protein-fractions are left-over flowthroughs after sequential DNA and RNA extraction on spin filters using the Qiagen AllPrep kit (cat no. 80004). The lysis of patient biopsies and DNA and RNA extraction was performed at Genomic Medicine (Copenhagen University Hospital, Denmark) [2]. They had had the foresight to keep and store the protein-fractions at-80°C from 2013 and onwards. For this study, we started with the protein-fraction flowthroughs and below describes all downstream processing in our laboratory. The final composition of the protein-fractions is difficult to assess. There will be guanidine salts (hydrochloride and thiocyanate), dithiothreitol (DTT), and varying amounts of ethanol present in each sample. Additionally, the procedures have gone through some minor changes since 2015, and there might hence be variations in sample compositions due to those that are hard to track.

### Pelleting of proteins

Advanced Protein Purification buffer (APP; Qiagen): the APP buffer contains zinc chloride to precipitate protein at room temperature. Following the manufacture’s instruction 1:1 volume of APP buffer to protein-fraction was added to the sample and mixed vigorously at room temperature. The solution was left for 10 minutes on the bench to precipitate the proteins. The solution was then centrifuged at 20,000 relative centrifugal force (RCF) for 10 minutes, and the supernatant was discarded. Left is a relatively large white pellet that was washed with 500 µL of 70 % ethanol and centrifuged at 20,000 RCF for 1 minute. Most of the ethanol was removed, and the rest was allowed to air-dry for 5 to 10 minutes on the bench.

Acetone precipitation: as an alternative to the APP buffer for protein precipitation, we performed standard acetone precipitation. Briefly, we added four volumes of ice-cold (-20°C) acetone (100 %) to each protein-fraction. The tubes were inverted a few times to ensure mixing and then incubated overnight at-20°C or-80°C. The next day a distinct cloudy precipitate had formed. Samples were centrifuged for 10 minutes at 20,000 RCF in a pre-cooled centrifuge at 4°C and the supernatant was discarded. The pellet was washed once with ice-cold (-20°C) acetone, using approximately 500 µL. The pellet was then air-dried for about 10 minutes, or until there was no more liquid above it, but without over-drying it.

### Resuspension of pellets

Pellets resulting from either APP or acetone precipitation were resuspended in several different buffers (from 50 to 500 µL), and with the help of vigorous pipetting and sonication (Bioruptor® Plus). Buffers used were: 1 % (w/v) Sodium Deoxycholate (SDC; PreOmics), or 1 to 4 % sodium dodecyl sulfate (SDS), or 50 mM 2-amino-2-(hydroxymethyl)propane-1,3-diol;hydrochloride (tris-HCl; pH 8.0) buffers, or 8M urea. Sonication was performed on a Bioruptor® Plus, with 15 cycles of 30 second on and 30 second off.

### Tryptic digestion and peptide cleanup using the Protifi S-trap™ kit

Protein extraction and peptide generation were performed using the Protifi S-Trap™ kit (Protifi) according to the manufacturer’s instructions. Briefly, an aliquot of the protein-fraction was reduced and alkylated with Protifi reducing and alkylating reagents. The reduced and alkylated proteins were then digested with trypsin using the Protifi digestion kit, and the resulting peptides were desalted using Protifi C_18_ spin columns. The desalted peptides were then eluted in 80 % acetonitrile (ACN) and dried down using a SpeedVac. The dried peptides were resuspended in 0.1 % formic acid for mass spectrometry analysis. It should be noted that the use of guanidine-HCl buffer is incompatible with the initial SDS-containing buffer provided in the Protifi kit, as the mixture becomes gelatinous. Therefore, the first step of the protocol that involves using the SDS-containing buffer was excluded.

### Tryptic digestion and peptide cleanup of APP or acetone protein pellets

After protein resuspension of protein pellets, either resulting from APP or acetone precipitation, and resuspended in either buffer (SDC, SDS, Tris, or urea), we reduced and alkylated and digested the protein using standard mass spectrometry proteomics protocols. Briefly, we have roughly 250 µL of each sample at this point and add 2.5 µL of 1M DTT and leave this at room temperature for 20 to 30 minutes. For alkylation we add 25 µL of 550 mM CAA and incubate in dark for 20 minutes. For digestion we first added 16 µL LysC (0.5 µg/µL) and incubated for 2 hours on a shaker at 800 rpm. Then we added 16 µL trypsin (0.5 µg/µL) and incubated overnight on a shaker at 800 rpm. The next day, digestion was stopped by acidification using formic acid to a final concentration of 1 %.

The resulting peptides were desalted on either styrene-divinylbenzene reverse-phase resin (SDB-RPS) StageTips or Sep-Pak C_18_ (1cc, 3mg, or 1cc, 3mg, or 1cc, 3mg; Waters), depending on the amount and volume. Briefly, for the SDB-RP stage tipping, 100 µL of 0.2 % trifluoroacetic acid (TFA) wash buffer was added to the StageTip, followed by the addition of 100 µL of the sample. The mixture was then centrifuged for 5 minutes at 750 RCF. The StageTip was then washed with 150 µL of 99 % isopropanol + 1 % TFA and centrifuged again for 5 minutes at 750 RCF. Afterward, the StageTip was washed with 150 µL of 0.2 % TFA wash buffer and centrifuged again for 5 minutes at 750 RCF. A total of 60 µL of elution buffer (960 µL ACN, 48 µL ammonia, and 192 µL H_2_O) was added and the mixture was centrifuged for 3 minutes at 500 RCF, and then transferred to an MS plate. The solution was then subjected to a 30 to 45 minutes speed-vacuumed at 45°C, followed by solvation in 15 µL of 3 % ACN and 0.1 % FA (A*). The mixture was then placed on a thermo shaker at 37°C and 1300 rpm for 10 minutes. Finally, the sample was ready for peptide concentration determination.

For desalting we also used Sep-Pak cartridges tC_18_ (1 cc 50 mg, or 3 cc 200 mg cartridges). For desalting on the 3 cc 200 mg cartridge then the cartridge was first conditioned with 2 mL of 100 % ACN, followed by 2 mL of 50 % ACN / 0.1 % FA. Next, it was equilibrated with 4 times 2 mL of 0.1 % TFA. The samples were then spun at the highest speed of 20,000 RCF for 10 minutes and loaded onto the cartridge. The cartridge was washed/desalted with 3 times 2 mL of 0.1 % TFA, followed by washing/desalting with 2 mL of 1 % FA to remove the TFA. The elution was performed with 2 times 1 mL of 50 % ACN / 0.1% FA. The eluate was then speed-vacuumed at 45°C until dry and resuspended in 50 µl of A*. Finally, the sample was ready for peptide concentration determination.

### Protein identification and protein and peptide concentration determinations

Coomassie Brilliant Blue staining: To better visualize the protein content of these protein-fractions, especially early in the process, we took an aliquot (before digestion) and submitted it to Coomassie Blue staining following standard protocols for protein denaturation and SDS-page electrophoresis. Briefly, the protein samples were denatured by heating at 95°C for 5 minutes in sample buffer consisting of 62.5 mM Tris-HCl pH 6.8, 2 % SDS, 10 % glycerol, 0.001 % bromophenol blue, and 5 % β-mercaptoethanol. The denatured protein samples were loaded onto a 12 % polyacrylamide gel and electrophoresed using a Bio-Rad Mini-PROTEAN® system at a constant voltage of 200 V until the dye front reached the bottom of the gel (between 30 and 45 min). The molecular weight of the protein samples was estimated using a PageRuler™ Plus prestained protein ladder as a standard. The separated protein bands were visualized by staining the gel with Coomassie Brilliant Blue R-250 for 1 hour and destaining with 30 % methanol and 10 % acetic acid until the background was clear.

Bicinchoninic acid (BCA) assay protein concentration determination: The Pierce™ BCA Protein Assay Kit (ThermoFisher Scientific, cat no. 23225) was used for protein quantification. The assay was performed according to the manufacturer’s instructions. Briefly, protein standards or samples were mixed with the BCA working reagent and incubated at 37°C for 30 minutes. The resulting purple color was measured at 562 nm using a spectrophotometer. Protein concentrations were determined by comparing the sample absorbance to a standard curve generated using known concentrations of BSA.

NanoDrop™ spectrophotometer analysis: The NanoDrop™ spectrophotometer (Thermo Scientific) analysis was performed to determine the concentration and purity of the peptide samples (there seemed to be an unidentified contaminant obstructing our upfront protein analyses, and we did not trust those results). Two µL of each sample was pipetted and the absorbance at 280 nm was measured and used to calculate concentrations. Dilution factors were considered when relevant for either approach.

### Phosphopeptide enrichment on an AssayMAP robot using immobilized metal affinity chromatography (IMAC) high-capacity Agilent AssayMAP Fe(III)-nitrilotriaceticacid (NTA) cartridges

Automated phosphopeptide enrichment experiments using AssayMAP Fe(III)-NTA cartridges were performed with the Agilent AssayMAP Phosphopeptide Enrichment v2.0 App, included in the Agilent AssayMAP Bravo Protein Sample Prep Workbench v2.0 software suite on the AssayMAP Bravo Protein Sample Prep Platform/robot [54]. Chemistry conditions were systematically evaluated and their effects on phosphopeptide enrichment were explained in the Results and Discussion section. Representative on-deck samples and reagents used for phosphopeptide enrichment are shown in Table 1. Tryptically digested, desalted, and lyophilized samples in 2-mL tubes were diluted to 0.5–5.0 μg/μL with 80 % acetonitrile (ACN)/0.1 % TFA. The samples were then manually transferred from tubes into polypropylene 96-well plates, and a default protocol was used for phosphopeptide enrichment of a-casein, unless otherwise indicated. Samples were eluted from AssayMAP Fe(III)-NTA cartridges into a PCR plate containing neat formic acid for immediate sample acidification. Finally, sample plates were sealed and stored in an autosampler at 4°C until resuspension in A*, and half the volume was injected on a mass spectrometer.

### Peptides preparation with KingFisherTM Flex system

The protein fractions were processed using the protein aggregation capture protocol [55] on a KingFisherTM Flex System (Thermo ScientificTM)30 with MagReSyn® Hydroxyl beads (ReSyn Biosciences). The storage solution from the hydroxyl beads was substituted with 70% ACN. KingFisher deep-well plates were primed with either 1 mL of 95% Acetonitrile (ACN) or 70% Ethanol (EtOH). Each sample was treated with 100 µL of a digestion solution (60 mM TEAB buffer), with Lys-C and trypsin at an enzyme-to-substrate ratio of 1:500, dispensed into the KingFisher plates. Following this, the samples were mixed with 100 mM Tris-buffer (pH8.0) to achieve a total volume of 300 µL in the KingFisher plates, with an additional infusion of ACN to reach a final volume percentage of 70%. Subsequently, beads were introduced into the samples at a protein to beads ratio of 1:2. To facilitate protein aggregation, a two-step mixing process ensued, involving 1-minute intervals of medium-speed mixing followed by a 10-minute resting phase. Subsequent washes were carried out in 2.5-minute increments at a gentle pace without detaching the beads from the magnet. Digestion proceeded through 100 cycles of agitation lasting 45 seconds each, interspersed with a 6-minute pause overnight at 37°C. Protease activity was quenched by acidification with trifluoroacetic acid (TFA) to a final volume percentage of 1%.

### Liquid chromatography setups

EASY nLC 1200 liquid chromatography: Two hundred (200) ng of tryptic peptides were loaded onto an EASY nLC 1200 liquid chromatography (Thermo Fisher Scientific) for proteomic analyses. For phosphoproteomic analyses we loaded half of everything from the IMAC enrichment after elution of enriched peptides. We used a 50-cm column with a 75-μm inner diameter (New Objective, USA), which had been packed in-house using ReproSil-Pur C18-AQ 1.9-μm silica beads (Dr Maisch GmbH, Germany) to transfer the peptide to a mass spectrometer. The 70-minute gradient looked as follows: 3 % (buffer B) to 19 % in 40 minutes, then 19 % to 41 % in 20 minutes, followed by a wash and analytical column equilibration. The 120-minute gradient looked as follows (for DIA): 5 % (buffer B) to 30 % in 9 5minutes, then 30 % to 60 % in 5 minutes, followed by a wash and analytical column equilibration. The 145-minute gradient looked as follows: 2 % (buffer B) to 25 % in 110 minutes, then 25 % to 40 % in 15 minutes, followed by a wash and analytical column equilibration. The column temperature was kept at 60°C using a Sonation Nanospray Flex ™ Column oven.

Evosep One liquid chromatographer: Following the manufacturer’s instructions, we loaded 200 ng of the digested peptides onto a disposable Evotip C18 trap column (Evosep Biosystems). Briefly, the Evotips were first wetted with 2-propanol and activated with 0.1 % formic acid in acetonitrile. They were then equilibrated with 0.1 % formic acid before being loaded onto the trap column using centrifugal force at 800 RCF for 1 minute. After loading, the Evotips were washed with 0.1 % formic acid, and 200 µL of 0.1 % formic acid was added on top of the disks to prevent drying. The mass spec-ready peptides were then transferred to a mass spectrometer using the Evosep One LC system (Evosep Biosystems [27]). The eluted peptides were separated using a standard preset gradient method on a 15-cm PepSep column (150 µm inner diameter; Evosep) packed with 1.5 μm Reprosil-Pur C18 beads (Dr Maisch GmbH, Germany). The column was used to separate the peptides over a standardized Evosep 44-minute gradient (with a total sample throughput of 30 SPD) using buffer A (0.1 % formic acid in mass spectrometry grade water) and buffer B (0.1 % formic acid in ACN. The column temperature was kept at 60°C using a Sonation Nanospray Flex™ Column oven.

### Mass spectrometer setups

Orbitrap Exploris 480 mass spectrometer in data-dependent acquisition mode: For data-dependent (DDA) analysis on a Thermo Scientific Exploris 480 mass spectrometer (Thermo Fisher Scientific), the mass spectrometer was operated in ‘top-12’ mode, in which MS spectra were collected in the Orbitrap mass analyzer (300-1,750 m/z range, 60,000 resolution) with an automatic gain control (AGC) target of 300 % and a maximum ion injection time of 60 ms. The most intense ions from the full scan were isolated with an isolation width of 1.3 m/z. After higher-energy collisional dissociation (HCD) at a normalized collision energy (NCE) of 25, MS/MS spectra were collected in the Orbitrap (15,000 resolution) with an AGC target of 80 % and a maximum ion injection time of 22 ms. Precursor dynamic exclusion was enabled with a duration of 30 s.

Trapped Ion Mobility Spectrometry (TIMS) quadrupole Time of Flight (TOF) mass spectrometer (Bruker timsTOF Pro or timsTOF SCP) in either data-dependent, or data-independent acquisition mode: The liquid chromatographer was coupled online to the hybrid TIMS quadrupole TOF mass spectrometer via a 50-cm in-house packed column, or a 15-cm Pepsep column – both described above. In data-dependent (dda)-PASEF mode [22], the MS1 mass range was 100 to 1,700 m/z, the ion mobility range was set to 1.6 Vs cm-2 and 0.6 Vs cm-2, and accumulation and ramp times were specified as 100 ms with 10 PASEF ramps, and an 0.4-minute exclusion time. Collision energy was set from 20 eV (0.6 Vs cm-2) to 59 eV at 1/K0 (1.6 Vs cm-2). The data-indepentent (dia)-PASEF acquisition covered an m/z-range of 100 to 1,700 at MS1 to 400-1,200 at MS2. The method included two ion mobility windows per diaPASEF scan with variable isolation window widths adjusted to precursor densities. Twenty-five diaPASEF scans were deployed at throughputs of 30 SPD (cycle time: 2.7 s). The ion mobility range was set to 1.6 Vs cm-2 and 0.6 Vs cm-2, and accumulation and ramp times were specified as 100 ms for all experiments. The collision energy was set from 20 eV (0.6 Vs cm-2) to 59 eV at 1/K0 (1.6 Vs cm-2). The original diaPASEF method was described previously [23, 24].

Orbitrap Exploris 480 mass spectrometer in data-independent acquisition mode for phosphoproteomics: The DIA method comprised one MS1 scan with a maximum injection time of 60 ms and AGC target was set to “Standard”, covering a range of 300 to 1,400m/z with a resolution of 120,000 for MS1 and MS2, respectively. The method also included 32 segments with varying isolation windows from 16 m/z to 157 m/z, a resolution of 30,000, and a maximum injection time of 54 ms for MS2. The stepped NCE was set at 25, 27.5, and 30.

### Proteomic and phosphoproteomic searches

MaxQuant for DDA analyses of proteomes: The DDA raw files were processed using MaxQuant [16] version 1.6.7. The Andromeda search engine [56] was employed for peptide and protein identification at a false discovery rate (FDR) below 1 %. The human UniProtKB database served as the forward database (see below for more details about the FASTA databases used), and the decoy search used the automatically generated reverse database. The enzyme specificity was set to ‘Trypsin/P,’ and ‘LysC’ with carbamidomethylation of cysteine as a fixed modification and oxidation of methionine, and acetyl (protein N-terminus) as variable modifications. The maximum missed cleavage sites was set to 1 and the minimum number of amino acids required for peptide identification was 7. Proteins that shared peptides were grouped together, and label-free protein quantification was performed using the MaxLFQ algorithm [57] with ‘match-between-runs’ (MBR; default settings) enabled, and a LFQ minimum ratio count set to 1. Proteins with only one razor or unique peptide and those identified as reverse hits, potential contaminants, or only by site-modification were filtered out.

DIA-NN (data-independent acquisition neural networks) searches for DIA analyses of proteomes: The DIA raw files for proteomes were analyzed using DIA-NN 1.8.0.1 [25]. Maximum mass accuracy tolerances set to 15 ppm for both MS1 and MS2 spectra. Trypsin/P, was set as protease, with 1 missed cleavage and a maximum of 2 variable modifications from ‘N-term M excision’, ‘C carbamidomethylation’, ‘Ox(M)’, and ‘Ac(N-term)’. Precursor lengths ranged from 7 to 30 amino acids and had a 2 or 3 charge. Proteomes were analyzed using MBR. Protein inference was turned off in DIA-NN. Proteotypic peptides were annotated using the ‘Reannotate’ option in DIA-NN with an on-the-fly generation of a spectral library used. The quantification mode was set to ‘Robust LC (high precision)’, and all other settings were kept default. To ensure the accuracy of the results, DIA-NN’s output was filtered at precursor q-value < 1 % and global protein q-value < 1 %, following previously published recommendations [58] and similarly to a previous diaPASEF workflow [59].

Spectronaut for DIA analyses of phosphoproteomes: For analysis of phosphoproteomes Spectronaut (v. 14, Biognosys AG, Schlieren, Switzerland) was employed using the ‘BGS Phospho PTM Workflow’. Fixed modification was ‘Carbamidomethylation of (C)’ and variable modifications were ‘Acetyl (Protein N-term)’, ‘Oxidation (M)’, and ‘Phospho (STY)’. A 1 % FDR cutoff was set at peptide-spectrum match, peptide and protein group levels. The protein q-value experiment and run wide cutoffs were set to 0.01 and 0.05, respectively. The dataset was subjected to analysis with a sparse q-value. In phosphoproteomics experiments, a PTM localization cutoff of 0.75 was set.

### Orbitrap Astral Spectronaut searches

Peptides were loaded onto Evotips Pure and measured with a data-independent acquisition (DIA) method. 200 ng of peptides were partially eluted from Evotips with <35% acetonitrile and analyzed with an Evosep One LC system (Evosep Biosystems) coupled online to an Orbitrap mass spectrometer (Orbitrap Astral, Thermo Fisher Scientific) [42, 43]. Eluted peptides were separated on an 8-cm-long PepSep column (150 µm inner diameter packed with 1.5 μm of Reprosil-Pur C18 beads (Dr Maisch)) in a standard preset gradient method (21 min, 60 SPD) with a stainless emitter (30 µm inner diameter). The mobile phases were 0.1% formic acid in liquid chromatography (LC)–MS-grade water (buffer A) and 0.1% formic acid in acetonitrile (buffer B). Data were acquired in DIA mode. Each acquisition cycle consisted of a survey scan at a resolution of 240,000 (normalized automatic gain control target (AGC) of 500% and a maximum injection time of 100 ms. Fragment ion scans were recorded with a maximum injection time of 5 ms and with 200 windows of 3Th scanning from 380 − 980 m/z. Higher-energy collisional dissociation (HCD) fragmentation was set to a normalized collision energy of 25%.

### Raw file processing and bioinformatic analyses

Raw files were analyzed with directDIA workflow in Spectronaut v.19 [60]. Default settings were used. Variable modifications were ‘Acetyl (Protein N-term)’ and ‘Oxidation (M)’. Data filtering was set to ‘Qvalue’. ‘Cross run normalization’ was enabled with the strategy of ‘local normalization’ based on rows with ‘Qvalue complete’. FDR was set to 1% at both the protein and peptide precursor levels. Raw data was searched against the human proteome reference database (Uniprot March 2023). The coefficient of variation (CV) was calculated on the raw intensity. Data analysis and visualization were done using Jupyter Notebook with visual studio code in the environment of Python 3.11.3.

### FASTA databases used

In this study, we utilized two FASTA files for the human proteome and phosphoproteome, namely UP000005640_9606.fasta (20,594 gene entries) and the additional human proteome file UP000005640_9606_additional.fasta. Both were downloaded 2019 from UniProtKB. The main file contains well-curated protein sequences for the human proteome, while the additional file provides supplemental sequences derived from proteogenomic studies and alternative splicing events [61].

### Statistical analyses

All data was log_2_ transformed and proteomics data was median-MAD (median absolute deviation) normalized into robust z-scores before statistical and bioinformatical analyses, unless else is stated. No imputation was performed on any of the data.

Morpheus for hierarchical clustering and heatmaps: For hierarchical clustering analysis, the data were normalized using log2 transformation and median-MAD normalized. The data was then imported into Morpheus (https://software.broadinstitute.org/morpheus/) for unsupervised hierarchical clustering analysis using the one minus Pearson correlation as the distance metric [62]. The results were visualized as a heat map where rows represented proteins and columns represented samples. The dendrogram of the heat map showed the clustering of samples based on protein expression levels. Morpheus is a freely available tool with the source code deposited here: https://github.com/cmap/morpheus.js.

pFind v. 3.0 for open searches of peptide modifications: Peptide identification and quantification were performed using pFind v. 3.0 (http://pfind.org/) based on pFind v. 2.0 [17]. MS raw data files were converted to pFind-compatible files using the pBuild tool with the default settings. The search parameters included trypsin as the enzyme, a precursor mass tolerance of 20 ppm, a fragment mass tolerance of 0.1 Da, and a maximum of two missed cleavages allowed. Carbamidomethylation of cysteine was set as a fixed modification, and oxidation of methionine and acetylation of the N-terminus were set as variable modifications. The FDR was controlled to less than 1 % at the peptide-spectrum match level using the decoy database strategy. The output files were filtered using a minimum peptide length of 6 and minimum ion score of 20. Peptide quantification was based on the extracted ion chromatogram (XIC) area of each identified peptide. The XIC was extracted with a 10 ppm mass tolerance and a 0.05 Da fragment ion tolerance using pParse, and the XIC area was calculated with the pQuant software [63]. The relative abundance of each protein was calculated by summing up the XIC areas of all its identified peptides. The protein quantification results were further normalized by the median ratio method to correct for variations in sample loading and instrument sensitivity.

The Clinical Knowledge Graph: the clinical knowledge graph (CKG) [18], is a python-based framework to overlay biological and clinical data onto graphs of omics data. The CKG is an open-source platform with over 20 million nodes and 220 million relationships that represent relevant experimental data, public databases, and literature. Statistical and machine learning algorithms were integrated into the CKG to accelerate the analysis and interpretation of proteomics workflows. Interactions and pathways can be calculated and visualized and was used for the patient with a HER2 amplified colon cancer. We first identified proteins with high changes in abundance (top 10 %) and with an upregulation at progression compared to baseline. Next, we used CKG to filter for proteins associated to colon and/or liver cancer (DOID:9256 and DOID:3571) and for proteins that act in a HER2 pathway (reactome). Lastly, the CKG was used to extract protein-protein interaction (PPI) information in order to generate a PPI network with cytoscape [64].

GraphPad Prism for statistical analyses: Statistical analyses were performed using GraphPad Prism (v. 9). Kaplan-Meier analysis was used to estimate survival probabilities, and chi-square analysis was used to test for associations between categorical variables. Graphs were generated using GraphPad Prism.

Gene-set enrichment analyses for biological evaluation: Raw gene expression data were processed using a single sample gene-set enrichment analysis (ssGSEA version 2.0), a freely available gene set enrichment analysis tool (https://github.com/broadinstitute/ssGSEA2.0) based on the original GSEA [65]. The gene-set collection used for ssGSEA were obtained from the Molecular Signatures Database (MSigDB v. 7.4) and we used the C8 database, which is a cell type signature gene-set that is curated from cluster markers identified in single-cell sequencing studies of human tissues (http://software.broadinstitute.org/gsea/msigdb/).

### Data availability

The mass spectrometry proteomics data have been deposited in the Proteomics Identifications Database (PRIDE) [66] with the dataset identifiers PXD041313 and PXD056337. Available data includes the 6 timeline samples (3 stored for 5 years and 3 for one year), and a larger cohort of 46 BRAF (V600E) mutated samples. Twenty-seven of which were used in this publication to assess throughput and proteomic depths, while an additional 18 (with matching baseline and progression biopsies) were used in a sister-publication about treatment resistance, and an additional 20 were used to evaluate baseline characteristics of BRAF mutated cancers for yet another sister-publication. The last “baseline cohort” have some overlap to the samples analyzed herein. All 46 were processed and measured as one batch for convenience, and uploaded to have the same PRIDE identifier, again, for convenience (both for the uploaders and the downloaders of data). In the second PRIDE identifier we have uploaded Astral data and phosphoenriched data.

