## Supplementary Figures for "In depth profiling of the cancer proteome from the flowthrough of standard RNA-preparation kits for precision oncology"

2 Current address: Novo Nordisk A/S, Maaloev, Denmark.

*Corresponding author

**Abstract:** Cancer is a highly heterogeneous disease, even within the same patient. Biopsies taken from different regions of a tumor may stand in stark molecular contrast to each other. Therefore, the ability to generate meaningful data from multiple platforms using the same biopsy is crucial for translating multi-omics characterizations into the clinic. However, it is generally a cumbersome and lengthy procedure to generate DNA, RNA and protein material from the same biopsy. The Qiagen AllPrep kit is an accessible, straightforward, and widely used kit in clinics worldwide to process biopsies and generate genomic and transcriptomic data from tumors. We aimed to determine if high-quality proteomics data could also be obtained from the remaining material. Here, we investigated procedures for generating deep and quantitatively accurate proteomic information in high throughput from Qiagen AllPrep flowthroughs. With a number of refinements, we obtain in excess of 10,000 quantified proteins, from 60 samples per day, achieving a substantial coverage of the total proteome. Additionally, we successfully characterize the tumors using phosphoproteomics. Combining a standard kit with in-depth proteomics will be an attractive approach for clinics seeking to implement multi-omics-based precision oncology.

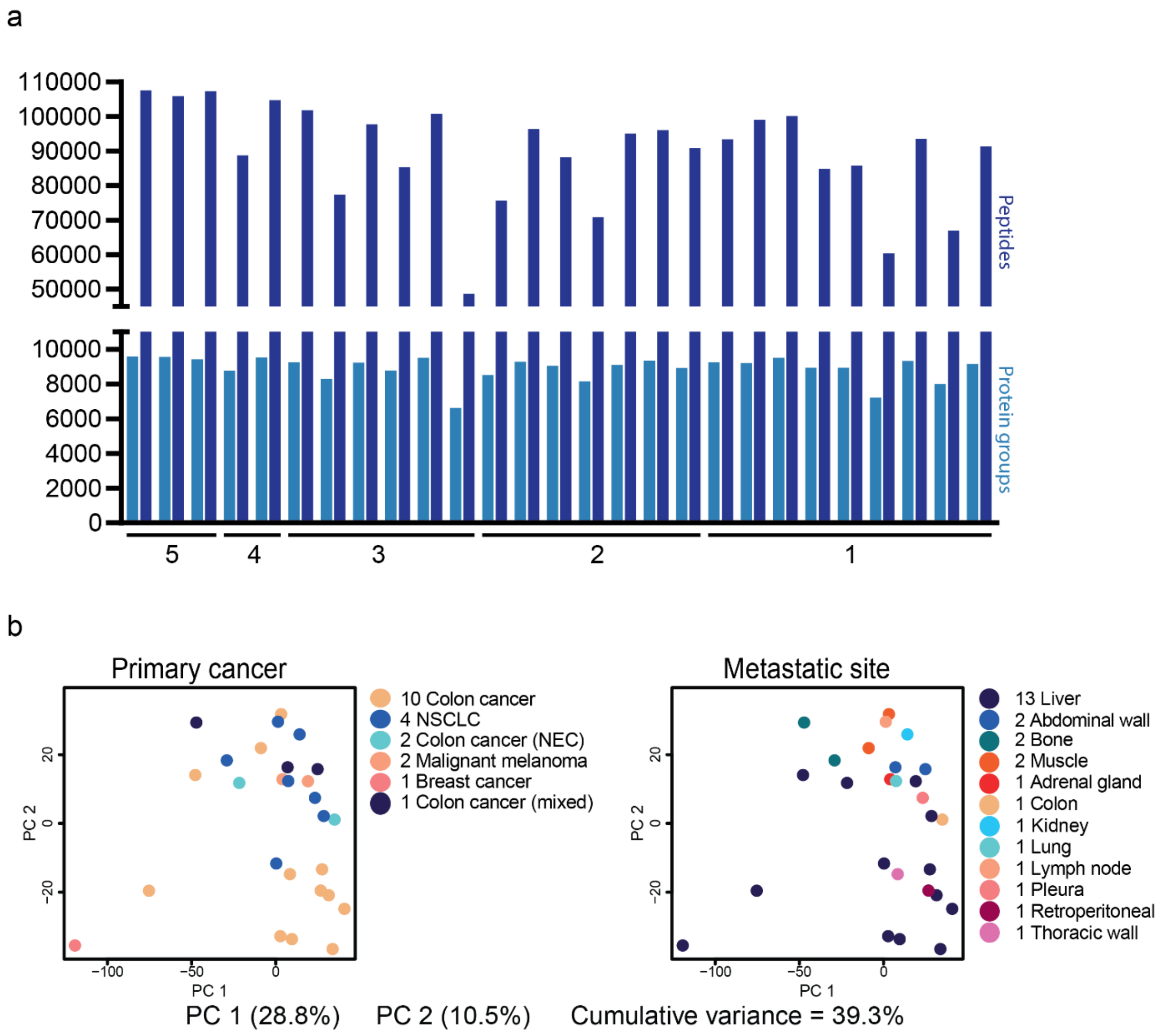


**Supplementary Figure S1. The 27 BRAF-mutated samples. a)** Numbers of quantified peptides and protein groups across all samples and for how many years they have been stored in -80°C (1 to 5 years). **b)** Principal Component Analyses (PCA) clustering of primary cancers and metastatic sites (in proteomic space). PC = principal component.

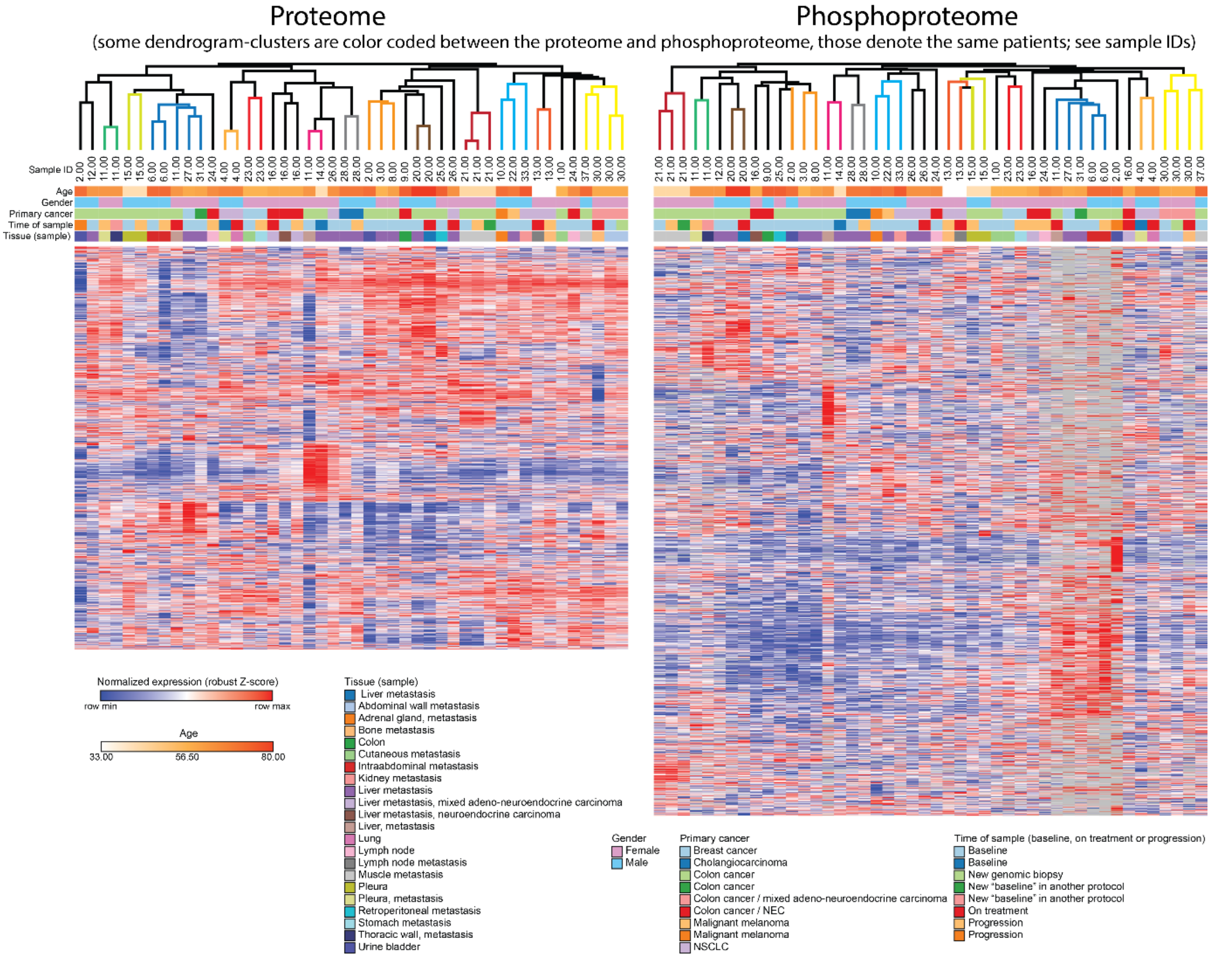


**Supplementary Figure S2. Unsupervised clustering of the 27 BRAF-mutated proteomes and phosphoproteomes.** One-minus Pearson clustering of samples in either proteome (left) or phosphoproteome (right) space. Note that the color coding of the dendrograms are matched, meaning that, for example, the yellow cluster (all the way to the left of both heatmaps) are the same patients coming together in both the proteome and in the phosphoproteome.
